# HEPN-AbiV is an RNase in the antiphage system AbiV

**DOI:** 10.1101/2024.05.05.592566

**Authors:** Xiaojun Zhu, Carlee Morency, Marie-Ève Picard, Cas Mosterd, Jason A. McAlister, Alice Perrault-Jolicoeur, Jennifer Geddes-McAlister, Rong Shi, Sylvain Moineau

## Abstract

Prokaryotes and eukaryotes possess defense systems, which can be either innate or acquired, to protect against viral infections. At the bacterial population level, abortive infection (Abi) serves as an innate immune defense mechanism against phage invasion. The AbiV antiviral system is prevalent in several bacterial genomes and exhibits diverse characteristics in terms of gene composition and evolution. Our investigation into the *Lactococcus* AbiV system revealed a novel two-component system, *abiV1* and *abiV2*, both of which are essential for its function as a type III toxin-antitoxin system. The toxin component AbiV (product of *abiV1*) is an RNase belonging to the HEPN (Higher Eukaryotes and Prokaryotes Nucleotide-binding) superfamily as it carries the consensus Rx4-6H motif. *In vivo* assays coupled with mass spectrometry showed that the lactococcal AbiV was expressed in the presence or absence of phages while *in vitro* experiments demonstrated that AbiV1 degraded ribosomal RNA but not mRNA. On the other hand, the antitoxin component (*abiV2*) was found to function as an RNA molecule that inhibited the nuclease activity of the AbiV1 toxin. The structural characterization of AbiV revealed that this RNase utilizes a large patch of positively charged area across the dimer to anchor RNA molecules. In addition, we showed that the AbiV N-terminal region (amino acids 1 to 23) is crucial for its RNase activity as a truncated AbiV lacking this segment adopted distinct conformational states incompatible with RNA binding. This study provided novel insights into the mode of action of the antiviral system AbiV.

## Introduction

Bacteriophages, or phages, are extremely abundant on our planet and their prevalence is usually higher than their bacterial hosts (1, 2). This vast quantity of phages plays a crucial role in maintaining the ecological balance. These phage-bacteria interactions also drive the coevolution of phages and their hosts (3, 4, 5).

To fend off viral attacks, bacteria have evolved multiple defense systems (6, 7, 8). There are now several types of antiviral systems that have been found in bacteria (9). Some of the most well-characterized systems enable the host cell to identify foreign genetic materials and cleave them. Examples of such mechanisms include CRISPR-Cas and restriction-modification (R-M) systems (10, 11). Programmed cell suicide mechanisms, on the other hand, involve cells killing themselves upon their detection of viral infection. Toxin-antitoxin (TA) and abortive infection systems are examples of such mechanisms (12).

Abortive infection, originally termed as non-productive phage replication, has been described in literature since the 1950s (13). A multitude of bacterial defense mechanisms collectively known as Abi have been discovered in various Gram-positive and Gram-negative bacteria, particularly in *Lactococcus* and *E. coli*. For example, over 20 types of Abi systems, named from AbiA to AbiZ based on their timing of discovery and characteristics, have been identified on plasmids and genomes of various *Lactococcus* strains (14). Most of the identified lactococcal Abi systems consist of a single gene, except for a few of them that encode multiple genes such as AbiE (15), AbiG (16), AbiL (17), AbiT (18), and AbiU (19).

Although many of these highly diverse lactococcal Abi systems were found decades ago and some of them have been even used by the cheese industry to improve the overall phage resistance of their starter cultures, the molecular mechanism has remained elusive for most of them. Abi systems typically act at various stages of the phage lytic cycle, and their activation depends on the nature of the system and the infecting phage. During the early stage of infection, Abi systems can interfere with phage DNA replication and RNA transcription (e.g., AbiA, AbiB) (20, 21). A recent study also demonstrated that the *Lactococcus* AbiA, which contains a HEPN domain, functions as a DNA polymerase in the AbiA antiphage system (22). Inhibition of protein synthesis is another strategy employed by lactococcal Abi systems, such as AbiC (23). Abi can also affect late gene expression (e.g., AbiK, AbiP) (24, 25, 26) and limit the packaging of phage virions (e.g., AbiI) (27).

Some Abi systems function through a toxin-antitoxin mechanism (28), which involves two molecules, such as AbiQ (29, 30) and AbiEi/AbiEii (31). Modulating the interactions between toxin and antitoxin will regulate the antiviral activities. In the type III TA-mediated AbiQ system, the toxicity of the protein AbiQ, which functions as an endoribonuclease, is neutralized by its repeated non-coding RNA antitoxin (30). Upon phage infection, AbiQ is activated to cleave cellular and phage RNA. In contrast, AbiE was found to be a type IV TA system (32). In the latter, the toxin protein AbiEii is predicted to be a nucleotidyltransferase while the antitoxin protein AbiEi contains a positively charged N-terminal winged helix-turn-helix domain that binds to two inverted repeats within viral promoters, leading to the transcriptional repression of phage genes.

The Abi phenotype can be obtained via different phage defense mechanisms (33). For example, Abi systems were associated with cyclic oligonucleotide-based antiphage signalling systems, collectively known as CBASS, found in various bacteria (34). The CBASS systems consist of an oligonucleotide cyclase and an effector protein. Upon phage infection, the oligonucleotide cyclase is activated, leading to the production of small messenger molecules known as cyclic oligonucleotides. These cyclic oligonucleotides, in turn, trigger the activity of a cell-killing effector protein. Depending on the system type, this lethal protein can be either a phospholipase or a membrane-spanning protein (35, 36).

Previously, we characterized the lactococcal abortive infection system AbiV, which is encoded on a bacterial genome and employs the protein AbiV to confer phage resistance (37, 38). Mutations in an early-transcribed phage gene, named *sav*, were also shown to confer insensitivity to AbiV (38). Others have later performed sequence-based analysis and identified a family of AbiV-like proteins, clustered within the HEPN superfamily, which is associated with a probable endoribonuclease activity (39). Here, we delved into the molecular mechanism of the lactococcal AbiV system. We reassessed its gene composition, used RNA sequencing and mass spectrometry to identify AbiV system transcription and expression, probed its RNase activity, and determined its crystal structure.

## Results

### AbiV is a HEPN-like protein found in diverse bacterial genomes

Protein Blast using the AbiV from strain *L. cremoris* MG1363 yielded 944 hits, showing that AbiV is present in many other bacterial genomes. A phylogenetic tree based on protein sequences revealed numerous branches, suggesting a significant evolutionary diversity within the AbiV cluster (Figure 1).

**Figure 1.**
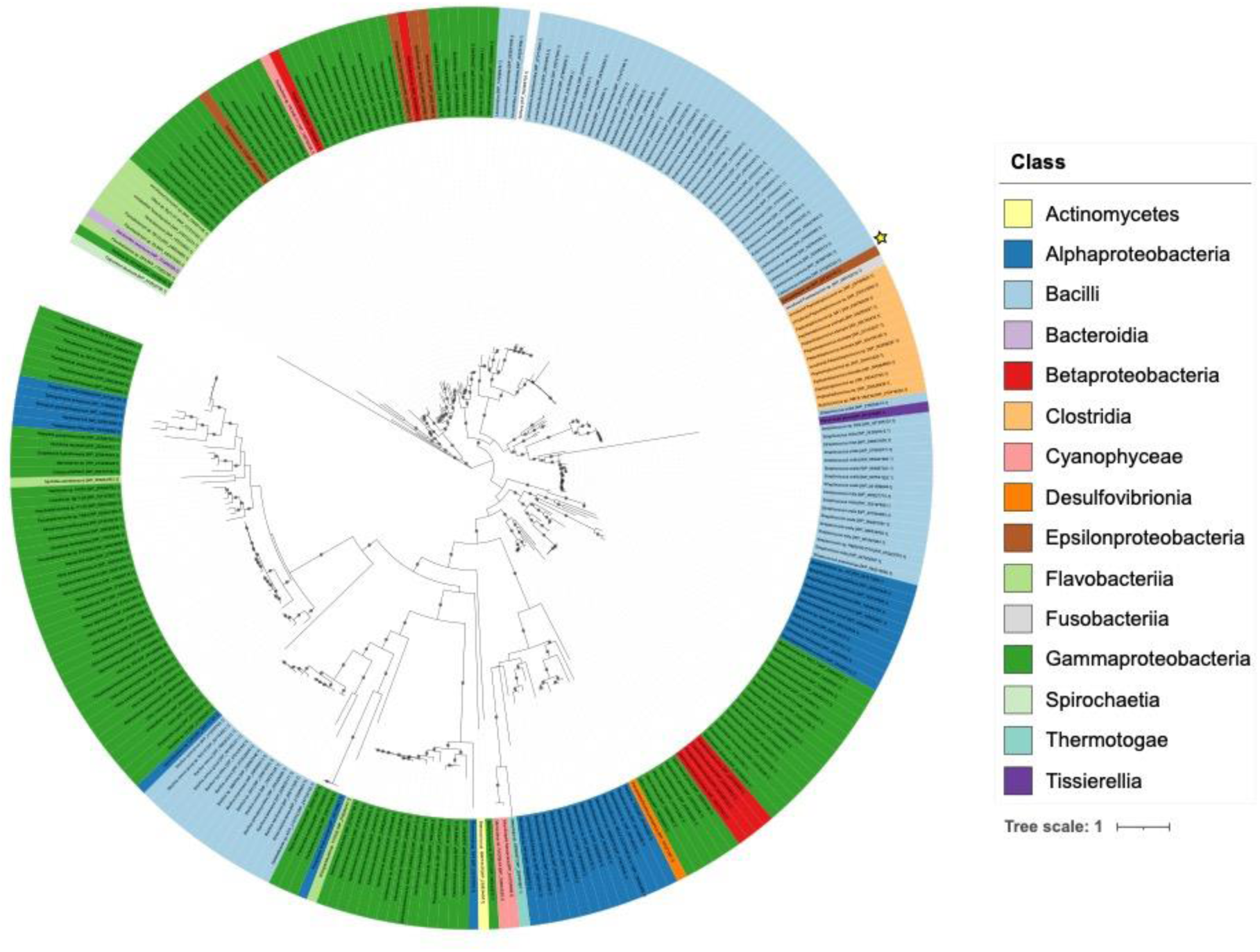
Phylogenetic analysis of the AbiV protein sequences. Branch support values greater than 80% are indicated by grey circles. Labels are color-coded based on bacterial class. AbiV from *L. cremoris* MG1363 is identified by a yellow star.

Moreover, it should be noted that the sequence identity within the AbiV family is not uniform (Supplementary Figure S1). Notably, a HEPN domain (30 aa to 170 aa) presumably responsible for nucleotide recognition, is highly conserved within the core region. Conversely, some motifs within this core region exhibited lower conservation. Also, the N-terminal region displayed size variability across AbiV-like proteins.

### Antiviral AbiV in *Lactococcus* is a two-component system

As AbiV alone was initially very difficult to express in *E. coli*, we suspected that it may be toxic. We reanalyzed the genetic architecture of the AbiV system and identified a putative small open reading frame (small *orf*, 180-bp) located upstream of *abiV* (named thereafter *abiV1)*, which may participate in the Abi system. First, we tried to clone a fragment containing only *abiV1* into the high copy vector pNZ123, the resulting plasmid was designated as *abiV1*::pNZ123 (Supplementary Table 1). Then, we cloned various regions flanking *abiV1* (Figure 2A, 2B), including with or without the small *orf*, to see their impacts on the AbiV-mediated antiviral activity. These plasmids were transformed into the laboratory phage-sensitive reference strain *L. cremoris* MG1363 to measure the anti-phage activity. The clone carrying only *abiV1* was transformed into MG1363, but only one clone out of three randomly selected clones had the intact *abiV* sequence. One clone exhibited a mutation leading to an amino change (P103T) in AbiV, while the other had a base deletion near its putative ribosome binding site. The other clones were readily obtained.

**Figure 2.**
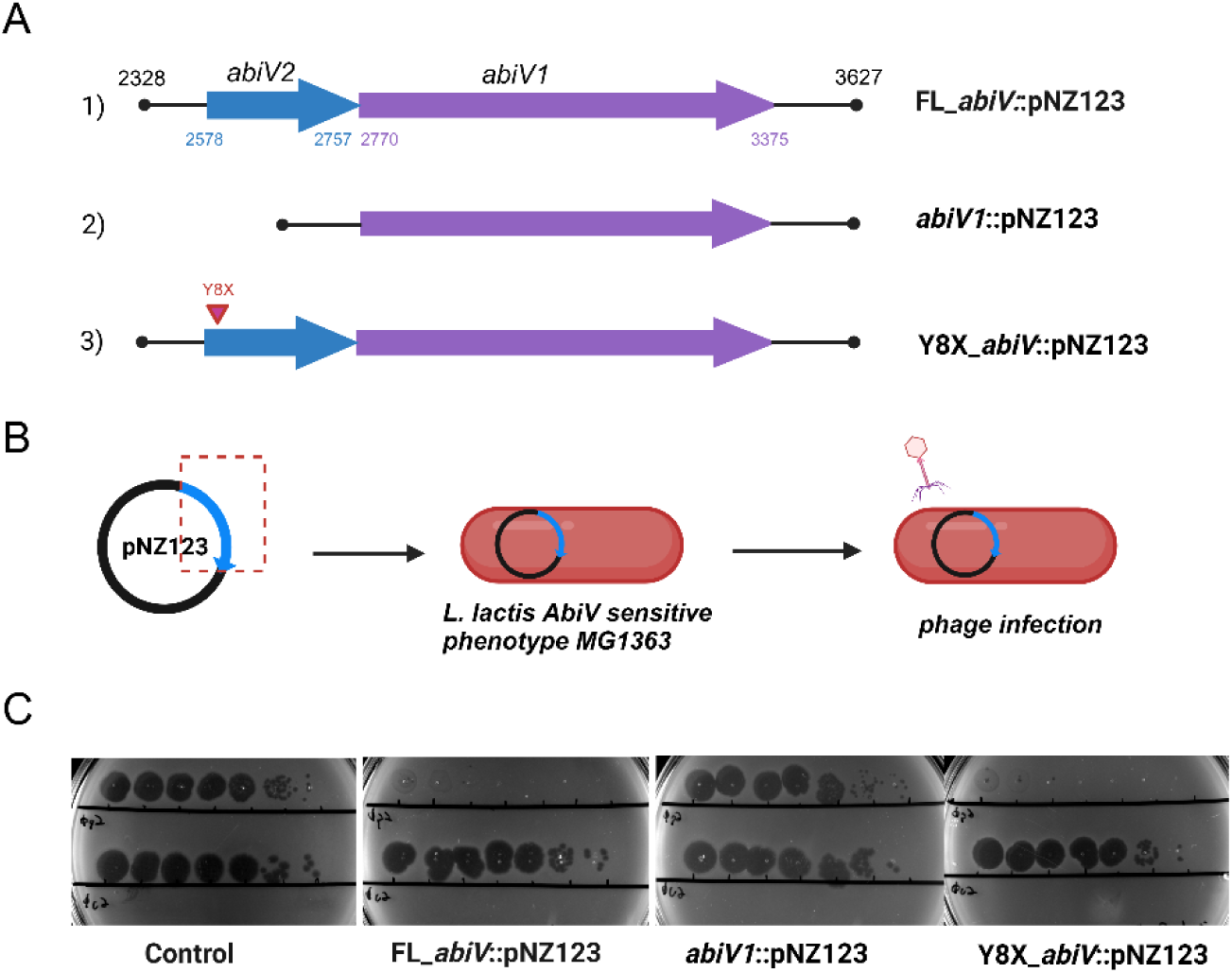
The construction of the AbiV antiviral system. Panel A 1) The full-length of the AbiV system, consisting of *abiV2* (position 698, 427 to 698, 596) and *abiV1* (position 697, 799 to 698, 404) from *L. cremoris* MG363 (GenBank AM406671.1) cloned into vector pNZ123. The numbers indicated the position in vector pNZ123. 2) Only *abiV1* was cloned into vector pNZ123. 3) The full-length of the AbiV system with a stop codon at Y8 in *abiV2*. Panel B) Cartoon illustrating that pNZ123 carrying the AbiV system and its derivatives was transformed into *L. cremoris* MG1363, and test for phage resistance. Panel C) Spot test assays of phages p2 (top) and c2 (bottom). *L. cremoris* MG1363 carrying an empty vector pNZ123 was used as a control.

*L. cremoris* MG1363 containing the *abiV1*::pNZ123 was sensitive to phage p2, while the derivative strain carrying instead the plasmid FL_*abiV*::pNZ123 led to a strong resistance to phage p2 (Figure 2C). On the other hand, the absence of the small *orf* resulted in a complete loss of the phage resistance phenotype (Figure 2C). This result indicated that the small ORF upstream of *abiV1* was essential for AbiV’s antiviral activity. Recognizing the significance of this putative small ORF, we designated it as “*abiV2*”.

### The AbiV system functions through an RNA-protein pair

*AbiV2* lacks a recognizable ribosome binding site and may initiate with an irregular start codon (TGG), suggesting that *abiV2* may not encode a protein, but instead may produce an RNA molecule. To verify this, we generated plasmid (Y8X_*abiV*::pNZ123) in which we introduced a stop codon at a putative tyrosine residue (8^th^ amino acid) to prevent translation of *abiV2* and tested for phage resistance as above. We observed that the *L. cremoris* strain carrying Y8X_*abiV*::pNZ123 still exhibited resistance to phage p2 (Figure 2C). This finding suggested that *abiV2* is likely acting as a non-coding RNA and work in concert with the protein AbiV to provide phage resistance.

### Transcripts of the AbiV system by RNA-sequencing

We investigated the transcription process of the AbiV system. We first used Cappable-seq and Term-seq RNA sequencing to determine the transcription start sites (TSS) and transcription termination sites (TTS), respectively. Analysis of the sequencing reads identified hotspots on FL_*abiV*::pNZ123. The putative TSS and putative TTS are summarized in Supplementary Table 2 and illustrated in Figure 3. The nearest putative TSS was found at 2558 and the nearest putative terminator at 2853. Although there is a clear peak in reads for this putative terminator site, there is also background noise surrounding this area. For *abiV1*, the nearest putative transcription start site was at 2617 while the nearest putative terminator was at 3553. Therefore, transcripts of *abiV2* and *abiV1* were predicted to be 296-nt and 937-nt long, respectively. Another possibility may be that *abiV2* and *abiV1* are co-transcribed as a longer transcript of 996-nt (2558–3553).

**Figure 3.**
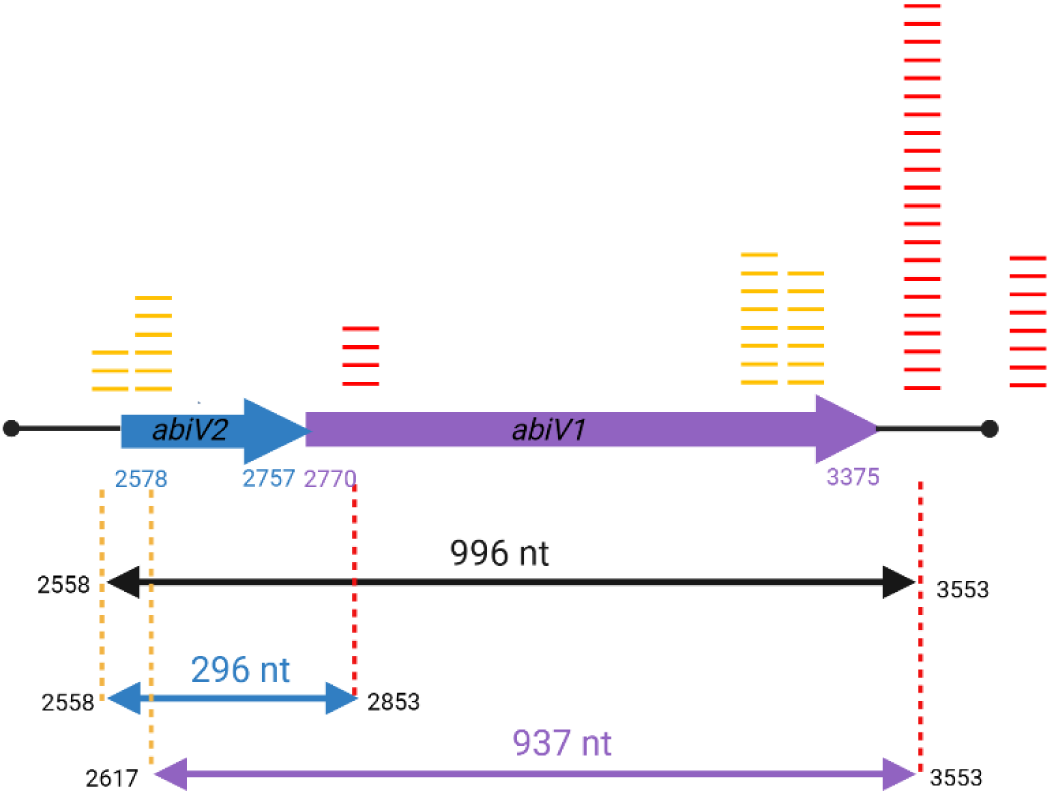
RNA-sequencing revealing the transcripts of the AbiV system. The sequence of AbiV in the vector pNZ123 is highlighted from 2328-bp to 3627-bp. *abiV2* spanning from 2578 to 2757 is flanked by *abiV1*, 2770 to 3375 illustrated by the large arrow. Transcription start sites are depicted as golden bars and terminators as red bars. Each bar represents 100 reads. The double arrow straight lines below indicate the putative transcripts length of *abiV2* (296 nt) and *abiV1* (937 nt). Two genes in the AbiV system may be transcribed to be a single long transcript of 996 nt, 2558 to 3553 (black arrow), or it may be transcribed separately to be a non-coding transcript of *abiV2*(2558–2853) and a coding *abiV1*(2617–3553).

### HEPN-AbiV cleaves ribosomal RNA *in vitro,* not mRNA

Many members of the HEPN family function as RNases (40). We performed an *in vitro* RNA degradation assay to check if AbiV is also an RNase. Our results indicated that AbiV degraded lactococcal 23S and 16S ribosomal RNAs, whereas two AbiV variants (AbiV_R91A and AbiV_R91A/H97A,) mutated in the HEPN catalytic motif, showed no rRNA degradation activity (Figure 4A). Furthermore, AbiV maintained its enzymatic activity in the presence of 10 mM EDTA, confirming that it is a metal-independent RNase. We then investigated whether AbiV has RNase activity towards mRNA. Considering the unknown role of *abiV2*, we used the 296-nt transcript of *abiV2* in the RNA degradation assay. *In vitro* RNase degradation assays revealed that the transcript of *abiV2* could not be cleaved by AbiV, although it can be efficiently digested by the broad-spectrum RNase A (Figure 4B). These results indicated that AbiV is an RNase and exhibited a preference for rRNA substrates.

**Figure 4.**
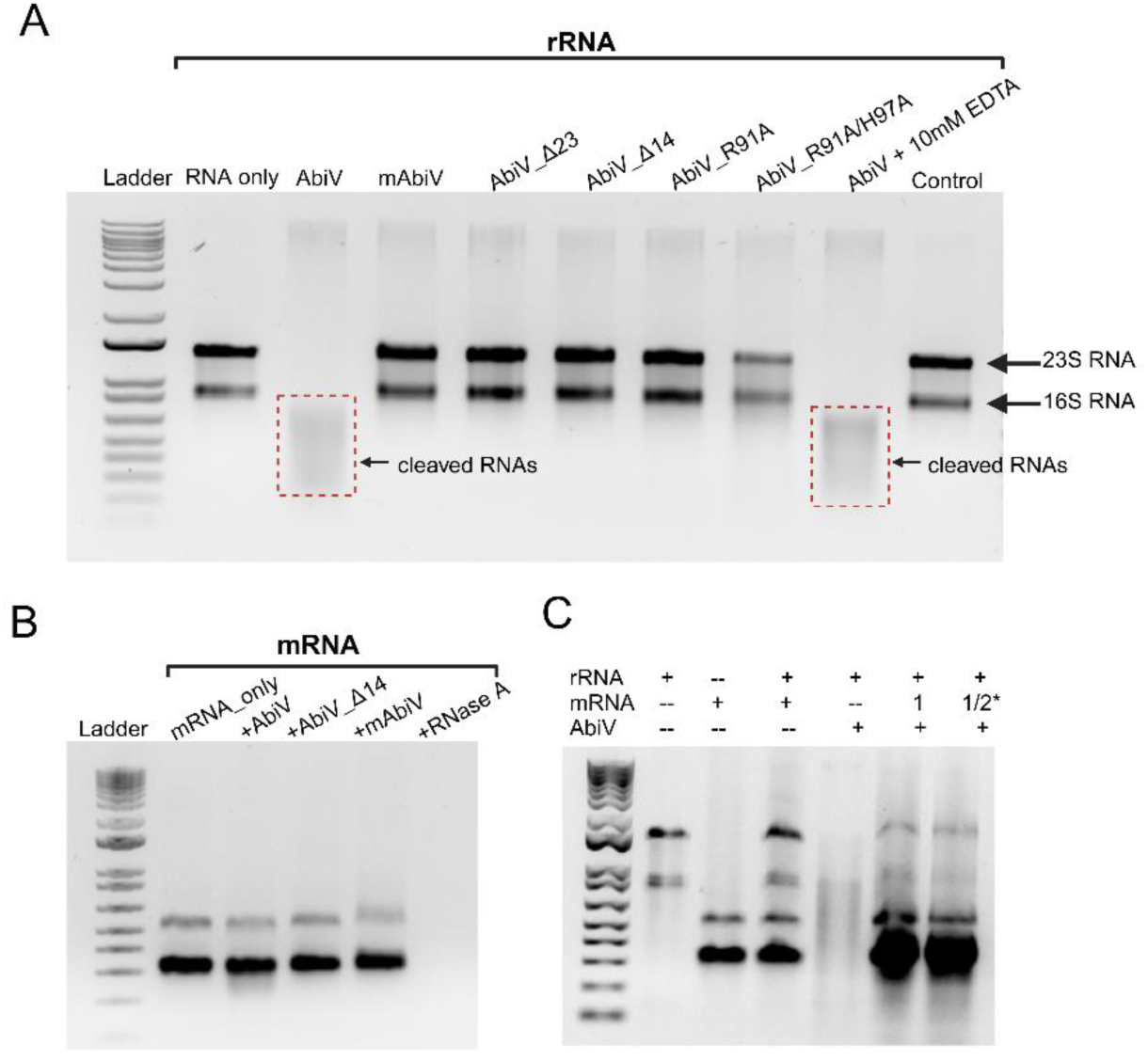
The *in vitro* RNA degradation and inhibition assay. Panel A) The wild-type AbiV degrades *Lactococcus* 23S and 16S rRNAs. The surface lysine methylated AbiV (mAbiV), N-terminal truncated AbiV (AbiV_Δ23, AbiV_Δ14), and the mutated variants in conserved Rx4-6H motif (AbiV_R91A, AbiV_R91A/H97A) cannot degrade rRNAs. The presence of 10 mM EDTA does not affect the RNase activity. Dashed zone indicates the degraded RNAs. The rRNA degradation reactions were run on 1% native agarose gel. Panel B) The 1.2% native agarose gel showed that AbiV1 is unable to degrade the *abiV2* mRNA transcript. RNase A was used as a positive control. The mRNA utilized in this assay was obtained by *in vitro* transcription using the RNA-sequencing map as a reference, spanning from 2558-to 2853-bp. Panel C) Transcript of *abiV2* inhibits the HEPN-RNase activity of AbiV against rRNAs. “1” indicates that mRNA of *abiV2* was incubation with purified protein at a molar ratio of 1.2 to 1. “1/2” indicates that the amount of mRNA incubated with AbiV was reduced in half.

### The transcript of *abiV2* inhibits the RNase activity of AbiV on rRNAs

As the AbiV system needs an RNA-protein pair, we explore the hypothesis that it may function as a type III toxin-antitoxin system, with *abiV2* RNA inhibiting the RNase activity of AbiV. The 296-nt RNA of *abiV2* was incubated with purified AbiV at a ratio of 1 to 1.2 and the resulting mixture was subjected to rRNA degradation assay. We observed that the presence of the *abiV2* transcript prevented the cleavage of rRNAs (Figure 4C). This finding suggests that *abiV2* RNA serves as an inhibitor of the rRNA degradation activity inherent to HEPN AbiV.

### AbiV selectively binds to a small hairpin RNA in transcript *abiV2*

The *in vitro* RNA degradation assay has demonstrated that transcript *abiV2* can block the RNase AbiV. Further investigation is needed to elucidate the interactions between AbiV and its neighboring RNA molecule, as well as the potential binding site. RNA molecules often adopt secondary structures crucial for function (41). Prior to conducting the RNA-binding investigation, the secondary structure of *abiV2* was predicted by the server MXfold2 (42) using RNA-sequencing-mapped sequence. The predicted model (Figure 5A) indicated a high likelihood of both paired and unpaired bases. To perform the AbiV-RNA binding assay, two RNA motifs of similar size (Figure 5B) were selected from different locations in *abiV2* in a random manner. It is worth noting that two selected RNA motifs are physically close. However, the gel mobility shift assay showed that only one of these motifs, nucleotides 85-to 102, interacted with AbiV (Figure 5C). This finding suggested that AbiV exhibits selective binding to RNA, particularly in the vicinity of the small hairpin motif.

**Figure 5:**
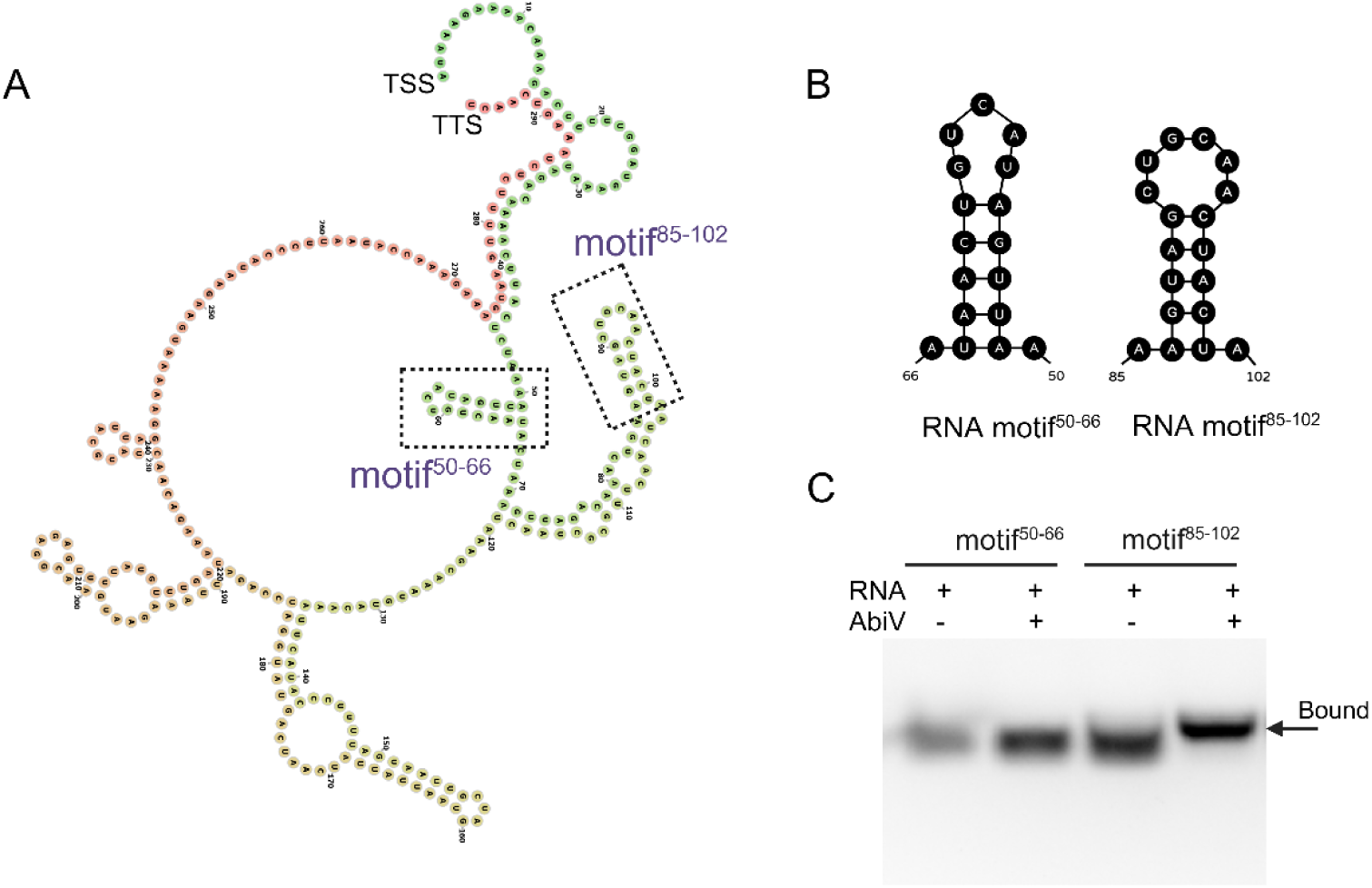
AbiV selectively binds to a small hairpin RNA motif. Panel A) The predicted secondary structure of the transcript *abiV2* as predicted by the MXfold2 algorithm. Base pairs are color-annotated from the transcription start site TSS (green) to transcription termination site TTS (pink). The small RNA motif^50-66^ and motif^85-102^ are highlighted within the overall structure. Panel B) Two RNA motifs containing hairpins with base details displayed. Both motifs feature a five-base-paired helix. Motif^50-66^ forms a five-base hairpin, while the other motif forms a six-base. Panel C) Gel mobility shift assay used to detect the AbiV-RNA interactions. RNA and protein were incubated at the molar ration of 2.5 to 1 at room temperature for the binding test.

### AbiV does not affect normal cell growth

In general, HEPN proteins exhibit toxicity within the cell (39). Although the free form of AbiV can degrade ribosomal RNAs, introducing the AbiV system into the cell did not affect cell growth. Therefore, we were motivated to explore its expression during regular cellular growth. To address this, we employed a mass spectrometry-based proteomics technique to detect the expression of AbiV in normal cell growth and at different timepoints during phage infection. The results obtained indicated that AbiV+ strains produced comparable levels of AbiV in the presence or absence of phage across a high-resolution time course (Figure 6). Conversely, AbiV was absent within the phage-sensitive strains in the presence or absence of phages (Figure 6). We could not detect AbiV2, supporting again that the small open frame in not translated.

**Figure 6.**
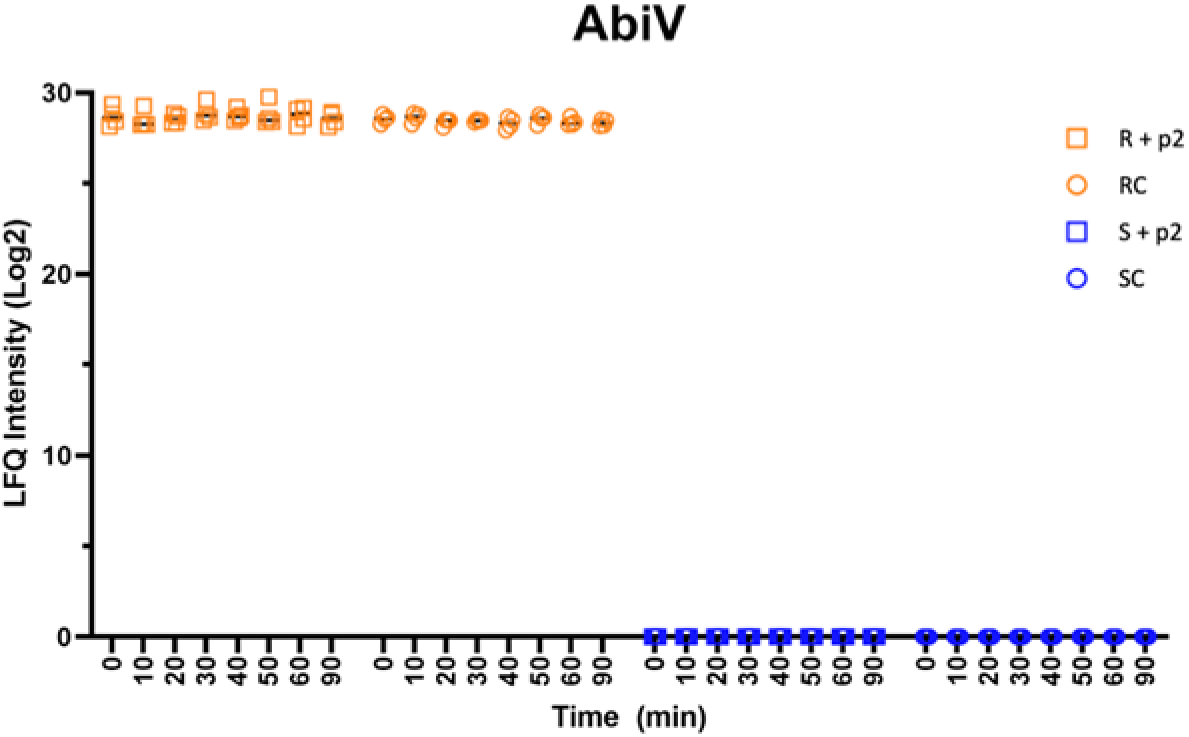
The expression levels of AbiV in phage-resistant and phage-sensitive *L. cremoris* strains. The phage-resistant strain *L. lactis* MG1363 carrying AbiV (FL_abiV::pNZ123) (R) produced comparable levels of AbiV1 (i.e., LFQ intensity) in the presence (R+P2, yellow square) or absence (RC, yellow circle) of phage p2 over time. In contrast, AbiV1 is absent in the phage-sensitive strain *L. lactis* MG1363 (pNZ123) in the presence (S+p2, blue square) or absence (SC, blue circle) of phage infection.

### Crystal structure of AbiV reveals a canonical HEPN dimer with notable flexibility

Crystallization of full-length AbiV was achieved only after a surface lysine methylation treatment. In both orthorhombic and monoclinic crystal forms, each asymmetric unit contained four AbiV subunits forming two biological dimers (Figure 7AB), like many other HEPN proteins (39, 43). In seven out of eight subunits in these two structures, a significant portion of AbiV, including the N-terminal region (amino acids 1 to 23) and a middle segment (aa 77-100), could not be reliably traced in the electron density maps (Figure 7C). In contrast, with the good electron density for the flexible regions mentioned above, most residues (aa 10-199) in the subunit A of the monoclinic crystal form could be successfully built (Figure S2AB). It should be pointed out that the Alphafold models of AbiV did not match well to the electron density map for the flexible region (Figure S2C). Excluding the above flexible regions, all the AbiV subunits had similar structures and could be superimposed with an r.m.s.d. of 0.4-0.6 Å for the ∼150 aligned Cα atoms. The most complete AbiV subunit was used for the description of secondary structure elements below. The AbiV subunit is composed of a core four-helix bundle (α2, α3, α7, and α8) surrounded by the one-turn helix α4, which is antiparallel to α7, and several other helical structures (α1, α5, and α6) stacking at different angles to the bundle (Figure 7A). The dimerization of AbiV was mainly achieved by the interactions from the structural elements α2, α3, loop α1/α3, and the β-hairpin segment (aa 141-151) of each subunit (Figure 7D). In particular, at least 10 H-bonds contributed by residues K33, D44, S53, T56, E67, K70, A142, and E154 could be identified in the dimerization interface with those incurred by E67 and K70 being the most conserved across the AbiV family.

**Figure 7.**
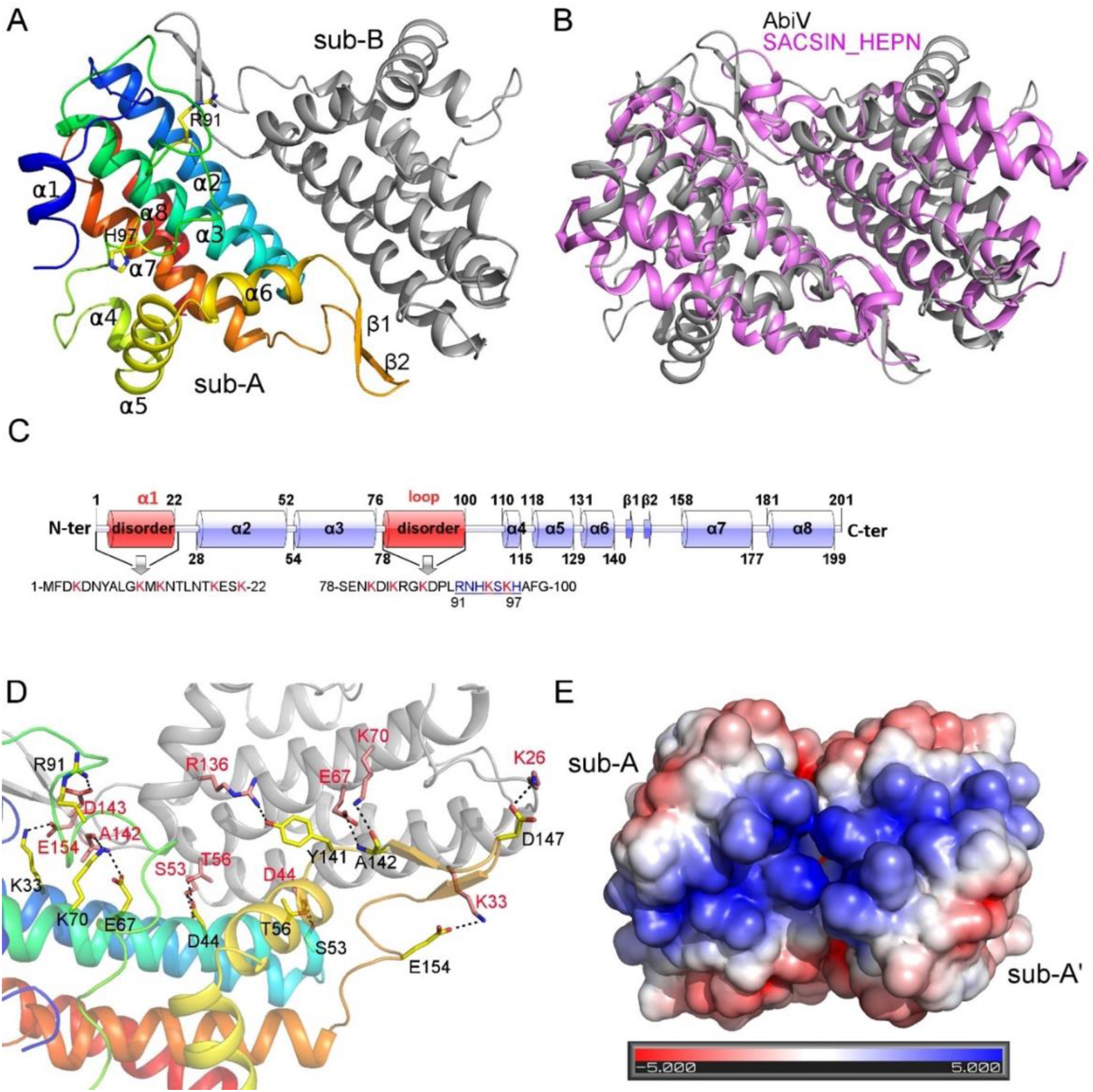
Crystal structure of AbiV showing a canonical HEPN dimer with a large positively charged patch for RNA binding. Panel A) Cartoon representation of AbiV dimer with sub-A painted in rainbow color (blue for N-terminal and red for C-terminal) and sub-B colored in gray. The secondary structures are labeled for sub-A with R91 and H97, the conserved residues in the RX_4-6_H motif (91-RNHKSKH-97) of the HEPN family, shown in stick mode. Panel B) Superposition of AbiV dimer (colored in gray) and the HEPN domain of SACSIN (PDB 3O10, colored in magenta). Panel C) A schematic diagram of the secondary structures in AbiV protein. The highly flexible regions (amino acids 1-22 and 78-100, colored in red) are largely disordered in 7 out of 8 independent subunits in the two crystal forms of AbiV. The sequences for these two segments are shown with lysine residues colored in red and the characteristic RX_4-6_H motif underlined. Panel D) The H-bonding interactions in the AbiV dimerization interface. The two subunits are colored in the same scheme as in A) and the residues forming the hydrogen bonds (shown in dotted lines) are shown in stick mode with their carbon atoms in sub-A and in sub-B shown in yellow and in salmon, respectively. Panel E) Surface representation of AbiV virtual dimer AÁ colored by electrostatic potential (ranging from −5 kBT/e (*red*) to 5 kBT/e (*blue*), calculated via PyMol (Schrödinger)) indicates that AbiV uses a large highly positively charged surface across the dimer for RNA binding. AbiV adopts the same orientation as in A). Due to the disordered two regions, the subunit B in the original AB dimer shown in A) is replaced by a more complete subunit A to better reflect the electrostatic potential of the dimer.

The Dali Server search revealed that the closest structural homologs of AbiV included the lincosamide antibiotic adenylyltransferase LinB (PDB 3JYY), the HEPN-domain containing protein AF0298 (2HSB), the TT1696 protein (1UFB), and the HEPN domain from human sacsin (43, 3O10) (Figure 7B), with a Z-score of 8.6-9.6 and an r.m.s.d. of 2.6-3.0 Å for ∼110 aligned Cα atoms, most of which were located on α2, α3, and α7. None of these closest structural homologs contain flexible regions as observed in AbiV structures.

### AbiV dimer contains a large positively charged patch for RNA binding

As a member of the HEPN superfamily, AbiV contains a conserved RX4-6H endoribonuclease motif. Inspection of the AbiV structure and the amino acid sequences of other AbiV-like members indicated that this short motif should be 91-RNHKSKH-97 (Figure 7C). The dimerization of AbiV led to the juxtaposition of two such catalytic subsets, allowing for nuclease activity upon RNA binding. Different from almost all other previously identified HEPN-domain containing proteins, this motif in AbiV was largely disordered in 7 out of 8 independent AbiV subunits. Nevertheless, the structure of amino acids 10 to 199 could be visualized in the subunit A of the monoclinic form (Figure S2B). Notably, while the imidazole ring of H97 had weak electron density, R91 in this subunit was clearly defined and stabilized via its H-bond with D143 of the other subunit in the dimer (Figure S2D). In this subunit, although the first ∼10 residues at the N-terminal could not be built, the other residues at the N-terminal (aa 10-22) formed the α1-helix and established favorable interactions with the α3-helix and the following loop α3/α4 as exemplified by the van der Waals contacts between M12 and L16 in α1 as well as I73, F76, and I77 in α3 as well as by the H-bonds formed between S21 in α1 and S78, N80, and D88 in the loop α3/α4 (Figure S2E).

To better reflect the electrostatic potential of dimeric AbiV, the subunit B with disordered regions (1-17 and 78-97) in the dimer was replaced by a copy of the more complete subunit A leading to a “virtual” dimer A-Á. As shown in Figure 5E, a large positively charged patch was formed across the dimer, indicating that the electrostatic interactions are likely the driving force to bind RNA. In the absence of RNA binding, the N-terminal region and the segment 78-97 were prone to be flexible as these are lysine-rich areas (K4, K11, K13, K19, K22, K81, K84, K87, K94, and K96) (Figure 7C). This may also explain why AbiV was recalcitrant to crystallization prior to lysine methylation. Notably, although the structure was from methylated AbiV, the electron density maps did not support the modeling of methylated side chains for most Lys residues and, in fact, many of the Lys side chains were still disordered (Figure S2D). Importantly, the crucial role of both the conserved HEPN motif 91-RNHKSKH-97 and the positively charged lysine residues were supported by the absence of RNA degradation activity with the AbiV variants R91A and R91A/H97A as well as by the methylation (loss of positive charges on AbiV1 surface) of surface lysine residues (Figure 7A).

### The N-terminal region of AbiV is critical for its structural integrity and RNase activity

In viewing of the disordered propensity of the N-terminal region of AbiV and its limited implication in the formation of a large positively charge patch of the dimer, we probed its role in the RNase activity. To achieve this, we constructed two shorter versions of AbiV through truncation of the first 14 or 23 residues (AbiV_Δ14 and AbiV_Δ23) and evaluated their RNase activities. Neither truncated AbiV retained any RNase activity, indicating that the N-terminal region is indispensable for its function.

Then, we crystallized AbiV_Δ23 and determined its structure. The N-terminal truncated AbiV was readily crystallized in the P1 space group and each asymmetric unit contained 24 subunits (sub_A to sub_X) assembling into 12 AbiV_Δ23 dimers (Figure S2F). The most remarkable feature of AbiV_Δ23 was its conformational diversity and various degrees of disorder of the 71 to 110 amino acids region in the different subunits. The most abundant conformation (conf_1) was present in 9 subunits (B, D, F, H, J, R, T, V, X) while the remaining 15 subunits could be divided into five other conformations (conf_2 to conf_6, 3 subunits for each), leading to four classes of AbiV_Δ23 dimers based on their subunit conformations: 6-1 for dimers A-B, C-D, and G-H; 5-1 for dimers E-F, Q-R, and S-T; 3-1 for dimers I-J, U-V, and W-X; 4-2 for dimers K-L, M-N, and O-P (Figure 8A). The conformation of the 71-110 region was the most complete in subunit K and drastically differed from that in the most complete subunit A of the full length AbiV (Figure 8B). Notably, although the HEPN motif 91-RNHKSKH-97 was well defined in both conf_3 and conf_6, this segment adopted completely different secondary structures, i.e., a loop in conf_3 and a helix in conf_6, which resulted in a displacement of 24.8 Å for the Cα atom of R91 upon superposition (Figure 8C). As a result of N-terminal truncation, the AbiV_Δ23 dimers did not seem to be compatible with efficient RNA binding. Taking the K-L dimer example (conf 4-2) as almost all the residues could be traced only in subunit K, we could replace the subunit L (76-92 being disordered) by a copy of subunit K leading to a virtual K-K′ dimer. Inspection of the electrostatic potential of this dimer revealed the absence of the positively charged surface usually found across the AbiV1 (Figure 8D). In fact, none of the four different AbiV_Δ23 dimers described above could unambiguously support the formation of the positively charged surface across the dimer. A plausible explanation is that the favorable interactions established between the N-terminal region (including α1) and the α3-helix as well as the following loop α3/α4 were important to maintain the overall conformation of the latter elements, which harbour a significant number of positively charge residues and the HEPN catalytic motif key for RNA binding and activity. In this regard, the N-terminal segment works as a “shoelace” to fasten the nearby structural elements preventing the latter from re-organizing into other distinct conformations incompatible for efficient RNA binding and/or cleavage.

**Figure 8.**
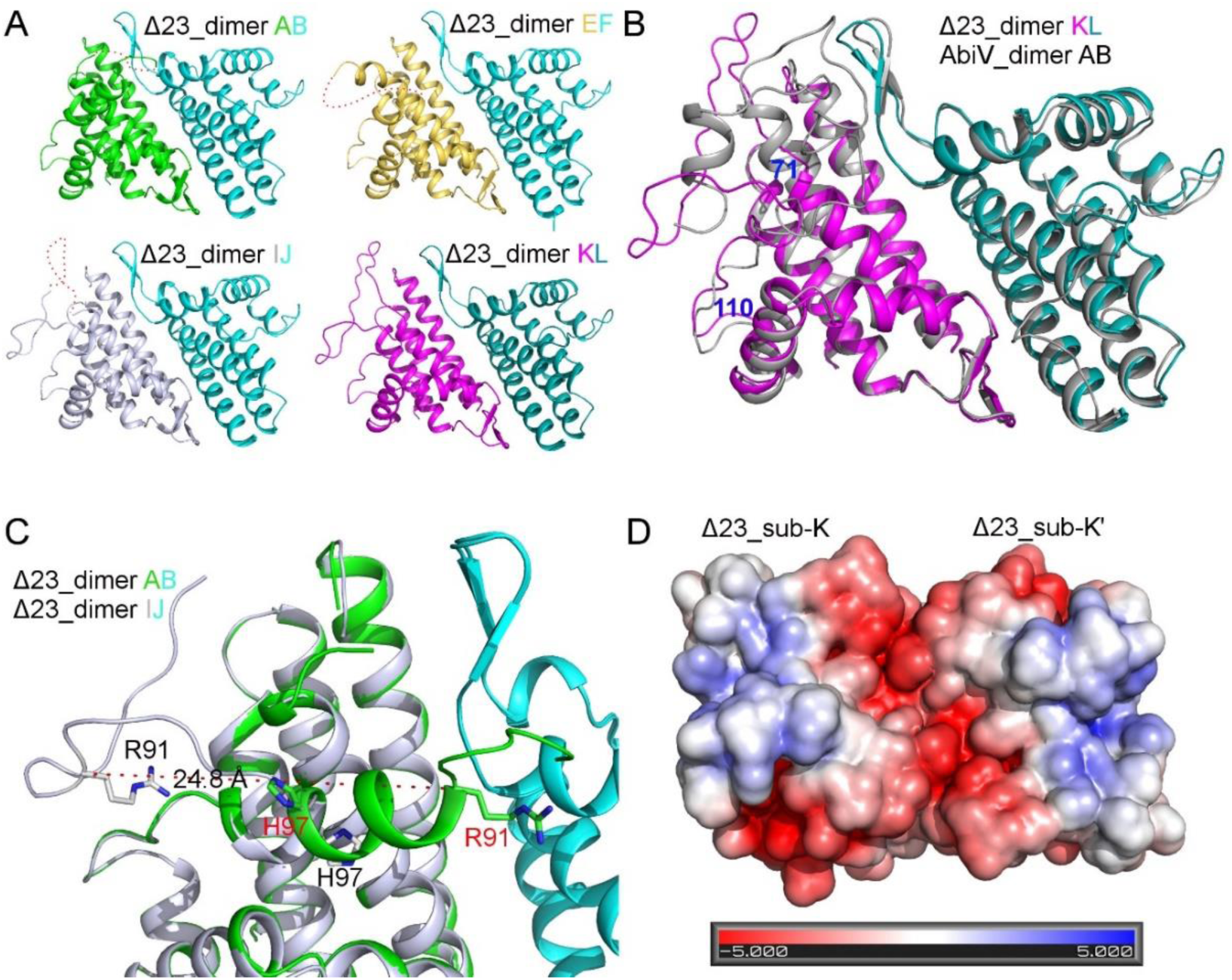
Crystal structure of AbiV_Δ23 revealing the importance of the N-terminal region in structural integrity required for RNA binding. Panel A) Conformational diversity of the amino acids 71 to 98 in different representative dimers and subunits of AbiV1. All six conformations (conf_1 to conf_6) are shown in cyan (B, F, J), teal (L), gray (I), magenta (K), yellow (E), and green (A), respectively. The unmodeled residues are indictated by red dotted curves. Panel B) Drastic conformational difference for the flexible amino acid segment 71 to 110 as shown in the subunit K (colored in magenta) of AbiV_Δ23 and in the subunit A (colored in gray) of full length AbiV1. Panel C) The characteristic HEPN motif (91-RNHKSKH-97) adopts distinct conformations, e.g., a helix in sub_A (colored in green) or a loop conformation in sub_I (colored in gray) in AbiV_Δ23. R91 and H97 are shown in stick mode with the Cα atoms of R91 being 24.8 Å apart (indicated by the dotted line) and those of H97 are in similar positions. Panel D) Surface representation of AbiV_Δ23 virtual dimer KK′ colored by electrostatic potential (ranging from −5 kBT/e (*red*) to 5 kBT/e (*blue*)) showing that AbiV_Δ23 loses the positively charged surface across the dimer for RNA binding due to conformational change upon N-terminal truncation. AbiV adopts the same orientation as in A). Due to the disorder segment 76 to 92, the subunit L in the original dimer KL shown in A) is replaced by a copy of the complete subunit K to better reflect the electrostatic potential of the AbiV_Δ23 dimer.

## Discussions

Bacteria possess a wide range of phage defense systems, including Abi systems, because they live in phage-containing ecosystems (44). Abi systems also function through varied modes of action, presumably to defend against a diverse phage population. In this study, we investigated the molecular mechanism of the lactococcal AbiV system through a multifaceted approach, which included gene composition, transcription and proteomic analyses, RNase activity assays, and structural characterization and comparison.

Our initial observations indicated that the AbiV protein was toxic to *E. coli*. Moreover, the transformation of *abiV1* alone led to the selection of clones with mutations. This prompted us to hypothesize that the AbiV system is likely part of a TA system. We showed that AbiV is an endoribonuclease. Notably, this enzyme possesses the capability to degrade host ribosomal RNAs, thereby suggesting a plausible mechanism for its cell-killing strategy during viral infections. Ribosomal RNAs are universally recognized as the prevailing type of RNAs in cells, serving as vital constituents of the ribosome, which plays a pivotal role in protein synthesis. Hence, degradation of host rRNAs would quickly lead to cell death and interruption of the lytic cycle upon viral infection.

Typically, in TA systems, the toxin is suppressed by an anti-toxin. Such anti-toxin was not originally identified in the previous characterization of the AbiV system (37). TA systems using protein-RNA pair as a toxin and antitoxin respectively usually belong to the type III (45, 46). In the lactococcal AbiQ defense system, another type III TA systems, the antitoxin is a small RNA which forms pseudoknots, directly binding the toxin via its unique gene architecture, a short palindromic repeat (30, 31, 47). However, we did not find such repeats within the vicinity of *abiV1*. However, through DNA-sequence and RNA-sequence analyses, we uncovered a small open reading frame (*abiV2*) with an intriguing transcription pattern, which included various transcripts. The introduction of a stop codon did not affect the phage resistance phenotype indicating that *abiV2* may act as an RNA molecule. In addition, while we could readily detect AbiV in *L. cremoris* cells (with and without phage infection), we could not detect AbiV2 under our conditions. The analyses of the transcription start sites (TSS) and transcription termination sites (TTS) suggested that the transcription of *abiV2* appears to be beyond its predicted DNA sequence, even encompassing segments of *abiV1*. In functional terms, unlike toxIN, the AbiV protein did not cleave the *abiV2* transcripts but its RNAse activity was inhibited by *abiV2*. Taken altogether, AbiV operates as a type III TA-like system. As no structure of AbiV was available, we determined the crystal structure of AbiV1, providing a prototype model for this family of proteins. The structural similarity between AbiV and other HEPN domain structures was limited to the core of helix bundle involved in dimerization (22). Despite its relatively small size (201 aa), AbiV differed from other HEPN structures previously determined as it exhibited a remarkable and unique flexibility for the amino acid segments 1-23 and 78-100. Indeed, the consensus RX4-6H catalytic motif was disordered in most subunits, which is usually not found in other structures containing a HEPN domain. Nevertheless, the only AbiV subunit, for which most residues (10–199) could be built, provided insights into the structural basis of AbiV to anchor RNA. The clustering of a series of Lysine, Arginine, and Histidine residues across the AbiV dimer contributed to a positive electrostatic potential to facilitate the binding of negatively charged RNA phosphate backbone. This is likely the underlying mechanism of many AbiV-like proteins to accommodate the RNA substrates for subsequent cleavage, as observed for the HEPN toxin SO_3166 (48).

The AbiV_Δ23 structure unexpectedly revealed multiple conformations of AbiV in one single crystal. In the absence of a largely disordered N-terminal region, the remaining part of AbiV could adopt drastically different conformations deviating away from the proper version required for functional RNase activity, revealing the molecular basis of inactive AbiV_Δ23.

Another notable feature of AbiV lied in the conformational plasticity of the HEPN RX4-6H catalytic motif, which was either disordered or well defined but in different secondary (loop or helical) structures. This structural characteristic may be unique as the RX4-6H motif was always located in a helical structure in other HEPN proteins (40). Moreover, such distinct conformational states of the RX4-6H motif were rarely observed in the apo form of other HEPN structures, although conformational changes or HEPN domain movements upon ligand/partner binding were commonly observed, as previously described for the Csm6’ ribonuclease (49) and the RNA-guided RNA ribonuclease Cas13a (50). This flexibility could also ease the structural adjustment of this motif for RNA binding and cleavage. As some of these conformational states were unlikely to be compatible for RNase activity, we could not exclude the possibility that, in addition to be inhibition by *abiV2*, the AbiV toxin could be further regulated by other cellular effectors via conformational modulation of the RX4-6H motif.

Although the 91-RNHKSKH-97 segment is the only region in AbiV that matches to the consensus RX4-6H motif in the HEPN family and that the R91A or R91A/H97A mutations abolished the RNAse activity of AbiV1, not all the members in the AbiV family contain such motif (39). The protein sequence alignment and our inspection of the Alphafold models (both monomer or dimer) for >100 AbiV family members indicated that neither R or H is strictly conserved and the spacing between these two residues may not be between 4 and 6, despite their highly conserved folding and dimerization pattern. Such divergences warrant further investigation on the distantly related members of the AbiV family.

In conclusion, we have investigated the mode of action of the AbiV system of *L. cremoris* MG1363. Our results established that the AbiV operates through a type III toxin-antitoxin like mechanism, comprising an RNA-protein pair that works collaboratively to prevent phage invasion. Moreover, our structural studies have provided molecular insights into the underlying mechanism of AbiV, an RNase that uses a large positively charged patch to anchor RNA molecules. In addition, we have also uncovered the structural basis of the crucial role of N-terminal region in the AbiV function. Nevertheless, further studies are required to better understand how the RNA of *abiV2* achieves its inhibitory activity on AbiV and how this toxin-antitoxin pair work concertedly to provide phage protection.

## Materials and Methods

### Phylogenetic analyses of AbiV

AbiV homologs were retrieved by protein blast homology 2.14.0 (51). Multiple alignment of AbiV-protein sequences was performed using MUSCLE 3.8.1551 (Edgar 2004). The alignment was used to generate a mid-rooted phylogenetic tree using IQ-TREE 2.2.1 (52) constructed with the General Q matrix (53) estimated from Pfam, version 31 (54), based on ModelFinder selection (55). Branch support was evaluated with 1000 ultrafast bootstraps from UFBoot2 (56). The tree was then exported and edited using iTOL, version 6.8 (57).

### Phage manipulations

All phages and bacterial strains used in this study are listed in Supplementary Table S1. Scrapings of phage lysates stocked in GM17 15% glycerol at -80°C were inoculated in 10 ml of growing host strain *L. cremoris* MG1363 in GM17 medium supplemented with 10 mM of CaCl_2_ and incubated at 30°C until clear lysis was observed. The cell lysate was centrifuged to remove bacterial debris and filtered through a 0.45 μm filter. A second phage propagation was carried out at 30°C by adding 50 μl of the first phage amplification to a 10 ml culture of the host strain when the OD_600nm_ reached between 0.1 and 0.2. Once a clear lysis was observed, the lysate was centrifuged and filtered (0.45 μm). The filtered lysates were stored at 4°C until use. Phage spot test assays were conducted as described previously (58). Phage p2 and c2 lysates were diluted 10-fold in buffer (50 mM Tris-HCl, pH 7.5, 100 mM NaCl, 8 mM MgSO4). Then, 500 µl of an overnight bacterial culture was mixed in molten soft agar (0.75%) at 55°C and poured on GM17 (Oxoid) plates supplemented with 1% agar + 10 mM CaCl_2_ + chloramphenicol (5 µg/ml). Last, 5 μl of phage dilutions were placed on the top layer of the plate and plates were incubated overnight at 30°C.

### Plasmid constructions

All plasmids and primers used in this study are listed in Supplementary Table S1. The main strategies used for plasmid constructions are as follows. To construct *abiV1*_only::pNZ123l, the primers *abiV1*_only_F and *abiV1*_only_R were first used to PCR-amplify *abiV1* from the genome of *L. cremoris* MG1363. The nucleotide positions of *abiV1* in *L. cremoris* MG1363 (GenBank AM406671.1) were 697, 765 to 698, 416. In parallel, the high copy vector pNZ123 was digested with XbaI and PCR-amplified using pNZ_XbaI_F and pNZ_XbaI_R2 primers (59). PCR products were purified using a Qiagen PCR purification kit and the linear pNZ123 was assembled with the *abiV* PCR product by Gibson assembly (60). The Gibson assembly product was dialysed and electroporated, into *L. cremoris* MG1363. The wild-type plasmid pNZ123 was also electroporated as a control in *L. cremoris* MG1363.

Colonies were selected on GM17 plates that contained chloramphenicol (5 µg/ml) and confirmed using pNZins_F and pNZins_R primers (59). To clone the complete AbiV system (positions 697, 547 to 698, 846) into pNZ123 (FL_*abiV*::pNZ123), the same method as the above *abiV* cloning was used except with the primers AbiV2_F and FL_*abiV* _R. To construct a full-length AbiV with a stop codon at Y8 into pNZ123 (*abiV2*Y8X_*abiV*::pNZ123), the primers Y8X_*abiV* _F and Y8X_*abiV* _R were used to amplify FL_*abiV*::pNZ123. The PCR product was purified using a Qiagen PCR purification kit and then phosphorylated using an Anza PNK kit (Invitrogen). Phosphorylation reactions were inactivated by incubating at 80°C for 5 min. Phosphorylated samples were ligated using a T4 DNA ligase kit (New England Biolabs) and incubated overnight at 16°C. The ligation product was electroporated into *L. cremoris* MG1363. Colonies were selected on GM17 plates with chloramphenicol and verified by PCR using pNZ123ins_F and pNZ123ins_R primers. All plasmid constructs were confirmed by Sanger sequencing.

### Protein expression and purification

The wild-type *abiV* was previously cloned in the expression vector pQE70 to generate pJH11, which was then introduced into *E. coli* M15, already containing pREF4, to obtain *E. coli* strain JH-62 (38). The strain JH-62 was grown overnight at 37°C, with agitation, in liquid LB medium supplemented with kanamycin 50 µg/ml (Kan50) to keep pREP4. The JH62 strain passed through a series of liquid medium passages without the ampicillin pressure, but with Kan50 to keep pREP4 and with an inoculation of 1% (61). After 12 passages, the culture was spread on LB + Kan50 plates and the colonies were streaked on LB + Kan50 and LB + Kan50 + Amp100 plates. The colonies that did not grow back on the antibiotic-containing plate were considered to have lost pJH11.

A plasmid was constructed (AbiV_Δ23::pQE70) to express AbiV that is deleted from its first 23 amino acids. Primers AbiV_Δ23_F and AbiV_Δ23_R were used to PCR amplify the region encompassing nucleotides 698, 401 to 698, 333 (AM406671.1) from pJH11. PCR products were purified, phosphorylated, and ligated. The ligation product was transformed into chemically competent JH62-delta-pJH11 cells. Colonies were selected on LB plates that contained Amp100 + Kan50 and verified by PCR using PQE70_F and PQE70_R primers. A similar plasmid (AbiV_Δ14:pQE70) was constructed as above but with AbiV deleted from its first 14 amino acids (nucleotides 698, 401 to 698, 360, AM406671.1). AbiV_R91A, and AbiV_R91A/H97A were obtained by gene synthesis. AbiV and variants were cloned into vector pQE70 with a C-terminal non-cleavable 6X His-tags.

The vector pQE70 carrying AbiV was transformed into expression *E. coli* strain M15. The recombinant *E. coli* cells were grown at 37°C in Terrific-Broth medium to reach an OD_600nm_ between 0.6 to 0.8. and protein overexpression was inducted with 1 mM isopropyl-beta-D-1-thiogalactopyranoside (IPTG). Cells were then cultured for another 16 hours at 18°C. Cell pellets were collected and resuspended in lysis buffer (50 mM Tris-HCl, pH 7.5, 300 mM NaCl, 5% glycerol, 5 mM imidazole, 1 mM PMSF) and lysed by sonication at 20 kHz after incubation with DNase (5 ug/ml) and lysozyme (2 mg/ml). Supernatants were collected by ultracentrifugation and then subjected to nickel affinity chromatography. After 2-step wash with imidazole-containing buffer (5 mM and 20 mM), the target protein was eluted with the buffer containing 200 mM imidazole. Eluted protein was then further purified by size exclusion chromatography using a Superdex 200 Increase (10/300GL) column with the sample buffer containing 20 mM Tris pH 7.3, 150 mM NaCl, and 1% glycerol. The N-terminal truncated AbiV (AbiV_ Δ23) was prepared using the same procedures. Lysine methylation of full-length AbiV was performed following the protocol described elsewhere (62).

### RNA extraction and sequencing

All equipment and reagents used were RNase-free whenever possible. *L. cremoris* MG1363 carrying FL_*abiV*::pNZ123 was grown until an OD_600nm_ of 0.6. Then, one ml of the culture was spun down, the cell pellet was re-suspended in 1 mL of TRIzol and incubated at room temperature for 5 minutes. A volume of 200 ul of chloroform was added and the sample was incubated at room temperature for 2 minutes, followed by centrifugation (12, 000 x g) at 4 °C for 15 minutes. The top fraction was mixed with 500 µL ice cold iso-propanol and incubated at room temperature for 10 minutes. The sample was centrifuged at 4 °C for 10 minutes and the pellet was washed three times in 75% ethanol. The pellet was re-suspended in water and treated with TURBO DNase (Thermo Fisher Scientific) according to the manufacturer’s instructions. The RNeasy MinElute Cleanup Kit (Qiagen) was used for further purification, after which RNA sequencing was performed by Vertis Biotechnology AG (Freising, Germany). Cappable-seq was performed to obtain the 5’ start site of the mRNA transcripts. Term-seq was performed to obtain the 3’ end of the transcripts. Sequencing was done on an Illuma NextSeq 500 system with 75 pb-length reads.

### *In vitro* RNA degradation and AbiV inhibition assay

rRNA molecules were obtained from *L. cremoris* MG1363 cultured using the above RNA extraction method. Total RNA was extracted and purified using the Qiagen RNA Mini Kit, and the quality of the RNA was measured by NanoDrop. Protein AbiV and inactive mutants were purified using the method above and the buffer changed to 25 mM Tris pH 7.3, 150 mM NaCl, 1% glycerol, and 5 mM MgCl_2_. mRNA was produced by *in vitro* transcription (Synbio Technologies). The sequence of mRNA used for the cleavage assay was determined by the map of RNA-sequencing, spanning from 2558 to 2853. For the *in vitro* RNase reaction, 0.2 µg of total RNA was incubated with 25 µg (final concentration 1.7 nM) of AbiV, surface lysine methylated AbiV, AbiV mutants, AbiV_ Δ23, or AbiV_ Δ14 at room temperature for 10 minutes. The reaction mixture was then loaded into a 1% native agarose gel and subjected to electrophoresis. For the mRNA cleavage assay, a 1.3% native agarose gel was used. To verify the inhibition of AbiV by mRNA, *abiV2* mRNA was incubated with AbiV at the molar ratio of 1.2 to 1 or 0.6 to 1. Then, the mixture of AbiV and mRNA was used in the rRNA degradation assay with the protocol described above.

### Crystallization, data collection, processing, and structure determination

The methylated full-length AbiV and the N-terminal truncated AbiV_ Δ23 were concentrated to 12 - 15 mg/ml in the sample buffer described above. Initial crystallization conditions were obtained using multiple crystallization screening kits from NeXtal Biotechnologies, following the method of microbatch-under-oil. Crystals were grown at room temperature, and subsequent optimization was performed to obtain crystals of diffraction quality. The crystals were flash-frozen in liquid nitrogen using the reservoir solution supplemented with 20% glycerol as the cryoprotectant before data collection at Canadian Light Source (Saskatoon, Canada) or at Advanced Photon Source (Chicago, USA). Methylated full-length AbiV yielded two different crystal forms: the first orthorhombic form was crystallized in the reservoir solution containing 0.1 M Tris-HCl pH 8.0, 1.8 M potassium sodium tartrate, and 0.4 M sodium bromide. and it was in the space group P2_1_2_1_2_1_ with the unit cell parameters of a = 73.69, b = 79.17, c = 136.14 Å. The second monoclinic form was obtained in the reservoir solution containing 1.0 M lithium chloride, 0.1 M sodium acetate, and 30% PEG-6000 and was in the P2_1_ space group with the unit cell parameters of a = 53.17, b = 144.96, c = 55.56 Å, β = 95.53°. The crystals of AbiV_ Δ23 were harvested from the reservoir solution containing 0.2 M lithium sulfate, 0.1 M Tris pH 8.5, and 30% PEG-4000 and belonged to the P1 space group with a = 74.44, b = 106.90, c = 146.23 Å, α = 90.96, β = 95.53, γ = 95.02°.

For phasing purpose, a one single-wavelength anomalous dispersion (SAD) dataset was collected at the wavelength of 0.91915 Å (Br-peak) for an orthorhombic crystal of methylated full-length AbiV obtained in the presence of 0.4 M sodium bromide, using the CMCF-BM beamline at the Canadian Light Source. The other datasets were collected at the wavelength of 0.97931 Å at the LRL-CAT beamline at the Advanced Photon Source of the Argonne National Laboratory. The SAD data of 2.55 Å were used for phasing via the AutoSol pipeline (63) and for initial model building via the AutoBuild program (64) in the Phenix package (65). The phases of both monoclinic crystal of full-length AbiV and the N-terminal truncated AbiV_ Δ23 were determined by MolRep (66) in the CCP4 suite (67) using the structure of the orthorhombic crystal as the search template. For all the reported structures, multiple cycles of refinement were carried out using Refmac5 (68) followed by model rebuilding with Coot (69). All the models have a reasonably good stereochemistry as analyzed with PROCHECK (70). Data collection and refinement statistics are shown in Table 1. The structures have been deposited with RCSB with the accession codes 9BJ5, 9BJ6 and 9BJ7.

**Table 1.**
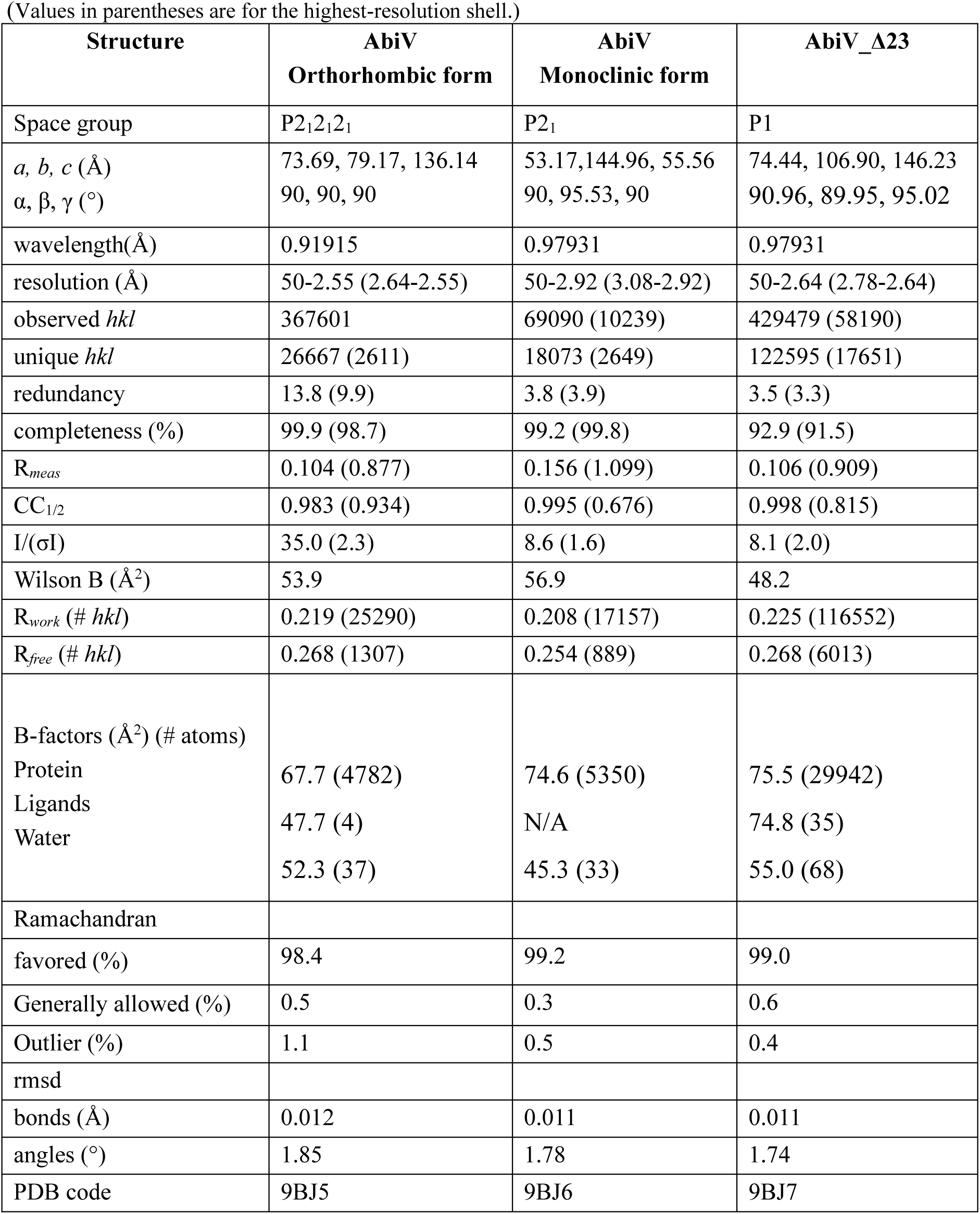
X-ray data collection and refinement statistics.

(Values in parentheses are for the highest-resolution shell.)
| Structure | AbiV<br>Orthorhombic form | AbiV<br>Monoclinic form | AbiV_Δ23 |
| --- | --- | --- | --- |
| Space group | P2<sub>1</sub>2<sub>1</sub>2<sub>1</sub> | P2<sub>1</sub> | P1 |
| *a*, *b*, *c* (Å)<br>*α*, *β*, *γ* (°) | 73.69, 79.17, 136.14<br>90, 90, 90 | 53.17, 144.96, 55.56<br>90, 95.53, 90 | 74.44, 106.90, 146.23<br>90.96, 89.95, 95.02 |
| wavelength(Å) | 0.91915 | 0.97931 | 0.97931 |
| resolution (Å) | 50-2.55 (2.64-2.55) | 50-2.92 (3.08-2.92) | 50-2.64 (2.78-2.64) |
| observed *hkl* | 367601 | 69090 (10239) | 429479 (58190) |
| unique *hkl* | 26667 (2611) | 18073 (2649) | 122595 (17651) |
| redundancy | 13.8 (9.9) | 3.8 (3.9) | 3.5 (3.3) |
| completeness (%) | 99.9 (98.7) | 99.2 (99.8) | 92.9 (91.5) |
| *R*<sub>meas</sub> | 0.104 (0.877) | 0.156 (1.099) | 0.106 (0.909) |
| CC<sub>1/2</sub> | 0.983 (0.934) | 0.995 (0.676) | 0.998 (0.815) |
| *I*/(σ*I*) | 35.0 (2.3) | 8.6 (1.6) | 8.1 (2.0) |
| Wilson B (Å<sup>2</sup>) | 53.9 | 56.9 | 48.2 |
| *R*<sub>work</sub> (# *hkl*) | 0.219 (25290) | 0.208 (17157) | 0.225 (116552) |
| *R*<sub>free</sub> (# *hkl*) | 0.268 (1307) | 0.254 (889) | 0.268 (6013) |
| B-factors (Å<sup>2</sup>) (# atoms) |  |  |  |
| Protein | 67.7 (4782) | 74.6 (5350) | 75.5 (29942) |
| Ligands | 47.7 (4) | N/A | 74.8 (35) |
| Water | 52.3 (37) | 45.3 (33) | 55.0 (68) |
| Ramachandran |  |  |  |
| favoured (%) | 98.4 | 99.2 | 99.0 |
| Generally allowed (%) | 0.5 | 0.3 | 0.6 |
| Outlier (%) | 1.1 | 0.5 | 0.4 |
| rmsd |  |  |  |
| bonds (Å) | 0.012 | 0.011 | 0.011 |
| angles (°) | 1.85 | 1.78 | 1.74 |
| PDB code | 9BJ5 | 9BJ6 | 9BJ7 |

### Time-course infection and proteomics sample preparation

The p2-sensitive *L. cremoris* MG1363 and its resistant derivative carrying FL_*abiV*::pN123 were pre-cultured in 10 ml of GM17 medium at 30°C and used to inoculate (1%) 100 ml of medium to grow until the OD_600nm_ reached 0.5. The culture was then centrifuged to remove the supernatant and the cell pellet was resuspended in 20 ml of fresh medium. A sample of uninfected bacteria (UI) was collected just prior to adding purified phage p2 at a multiplicity of infection (MOI) of 10. The sample T0 was collected immediately once p2 was added to the bacterial culture. The cultures were incubated at 30°C and samples collected at 10-, 20-, 30-, 40-, 60-, and 90-minutes post-infection, respectively. Protein extractions were performed as previously described (71). Briefly, *L. cremoris* samples were resuspended in 100 mM Tris-HCl (pH 8.5) with a protease inhibitor cocktail tablet (Roche) and 2% sodium dodecyl sulfate. Samples were homogenized by probe sonication (5 cycles of 30s on and 30s off; 30% power) in an ice bath followed by addition of dithiothreitol (10 mM final) and incubation at 95 °C for 10 min with shaking at 800 rpm. Samples were cooled and iodoacetamide (55 mM final concentration) was added followed by incubation at room temperature for 20 min (in the dark) and precipitation at -20 °C (overnight) in acetone (100%). Samples were pelleted, washed (80% acetone), resolubilized (8 M urea, 40 mM HEPES), quantified, and digested overnight (LysC and trypsin protease mix; protein:enzyme ratio of 50:1). Peptides were purified using STop And Go Extraction tips (STAGE-tips) (72), and dried in a SpeedVac prior to measurements.

### Mass spectrometry data processing and analysis

Samples were analyzed on an Easy-nLC 1200 high performance liquid chromatography system (ThermoFisher Scientific) coupled online to an Exploris 240 mass spectrometer (ThermoFisher Scientific). Samples were loaded onto an in-line 75-µm by 50-cm PepMap RSLC EASY-Spray column filled with 2 µm C18 reverse-phase silica beads and directly electrosprayed into the mass spectrometer using a linear elution gradient from 3% to 20% buffer B (80% ACN, 0.5% acetic acid) over a 3 h gradient at a constant flow of 250-nL/min. The mass spectrometer was operated in data-dependent mode with full scans of 400 to 2, 000 m/z acquired with a resolution of 120, 000 at 200 m/z. Raw files were analyzed using FragPipe software (version 19.0.) (73, 74, 75). The derived peak list was searched with MSFragger (73) against *L. cremoris* (Feb. 2023; 2383 sequences) and phage p2 (Feb. 2023; 49 sequences). The parameters were as follows: trypsin specificity (up to two missed cleavages), minimum peptide length (seven amino acids), fixed modification: carbamidomethylation, variable modifications: N-acetylation and oxidation, minimum of two peptides per protein, and peptide spectral matches and protein identifications were filtered using a target-decoy approach at a false discovery rate (FDR) of 5%. Relative label-free quantification (LFQ) and match between runs were used with a minimum ratio count of 1 (75). Protein filtering was performed using Perseus (version 1.6.2.2) (76). Proteins aligning to contaminants were removed, followed by transforming values to a log scale (log2), and valid-value filtering for three of four replicates in at least one group. Missing values were imputed from a normal distribution (downshift of 1.8 standard deviations and a width of 0.3 standard deviations). Data visualization utilized GraphPad Prism (version10.1.0, GraphPad Software) with the mean and standard deviation presented across four biological replicates at each time point.

### Data availability

The coordinates and structure factors of AbiV and AbiV_ Δ23 have been deposited in the Protein Data Bank with the accession codes 9BJ5, 9BJ6, and 9BJ7, respectively.

## Acknowledgements

We thank Simon Labrie for help with analyzing the RNA sequencing results. X.Z. was supported by the merit scholarship program for foreign students (PBEEE), from the Fonds de Recherche du Québec - Nature et Technologies (FRQNT), and by PROTEO. C.M. was also supported by a graduate scholarship from FRQNT. This project was supported by a FRQNT team grant to S.M and R.S. as well as by grants from the Natural Sciences and Engineering Research Council (NSERC) to R.S. (Discovery program) and S.M. (Alliance Missions). S.M. holds a T1 Canada Research Chair in Bacteriophages. Part of the research described in this paper was performed using beamline CMCF-BM at the Canadian Light Source, a national research facility of the University of Saskatchewan, which is supported by the Canada Foundation for Innovation, NSERC, National Research Council, Canadian Institutes of Health Research, Government of Saskatchewan, and University of Saskatchewan. This research used resources of the Advanced Photon Source; a U.S. Department of Energy (DOE) Office of Science User Facility operated for the DOE Office of Science by Argonne National Laboratory under Contract No. DE-AC02-06CH11357. Use of the Lilly Research Laboratories Collaborative Access Team (LRL-CAT) beamline at Sector 31 of the Advanced Photon Source was provided by Eli Lilly and Company, which operates the facility.

