## Supplementary material for "HEPN-AbiV is an RNase in the antiphage system AbiV": SuppMat_AbiV_XZ.pdf

### Supplementary Figures and Tables, Zhu *et al.*

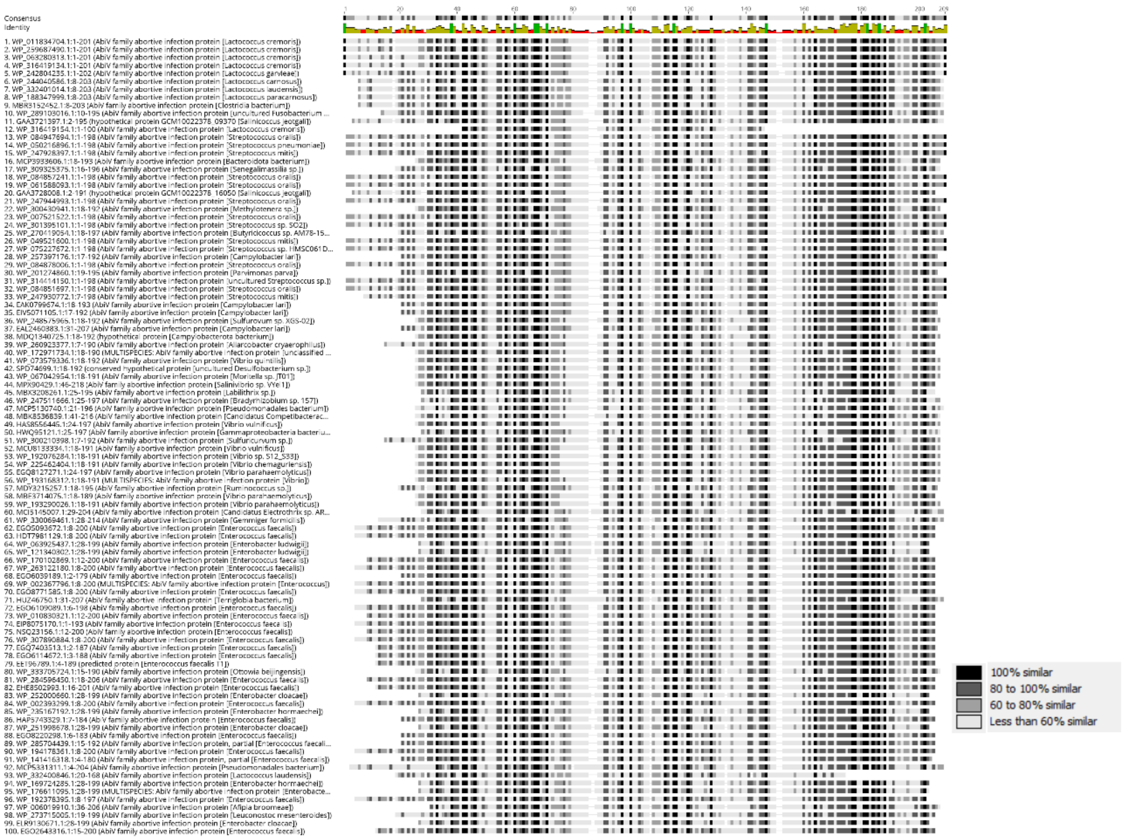


**Supplementary Figure 1: Sequence-based alignment of the AbiV family.** AbiV diversity analysis by aligning the top 100 homologs with amino acid identities ranging from 100% to 38.5% using Protein Blast. The region from positions 30 to 170 corresponds to the recognition region of the HEPN-AbiV superfamily. The red bars at the top of the figure indicate highly conserved regions. The red dashed borders highlight the variability in length across the N-terminal region of this superfamily protein. This figure was generated by *Geneious version 11.0 created by Biomatters. Available from* [*https://www.geneious.com*](https://www.geneious.com)

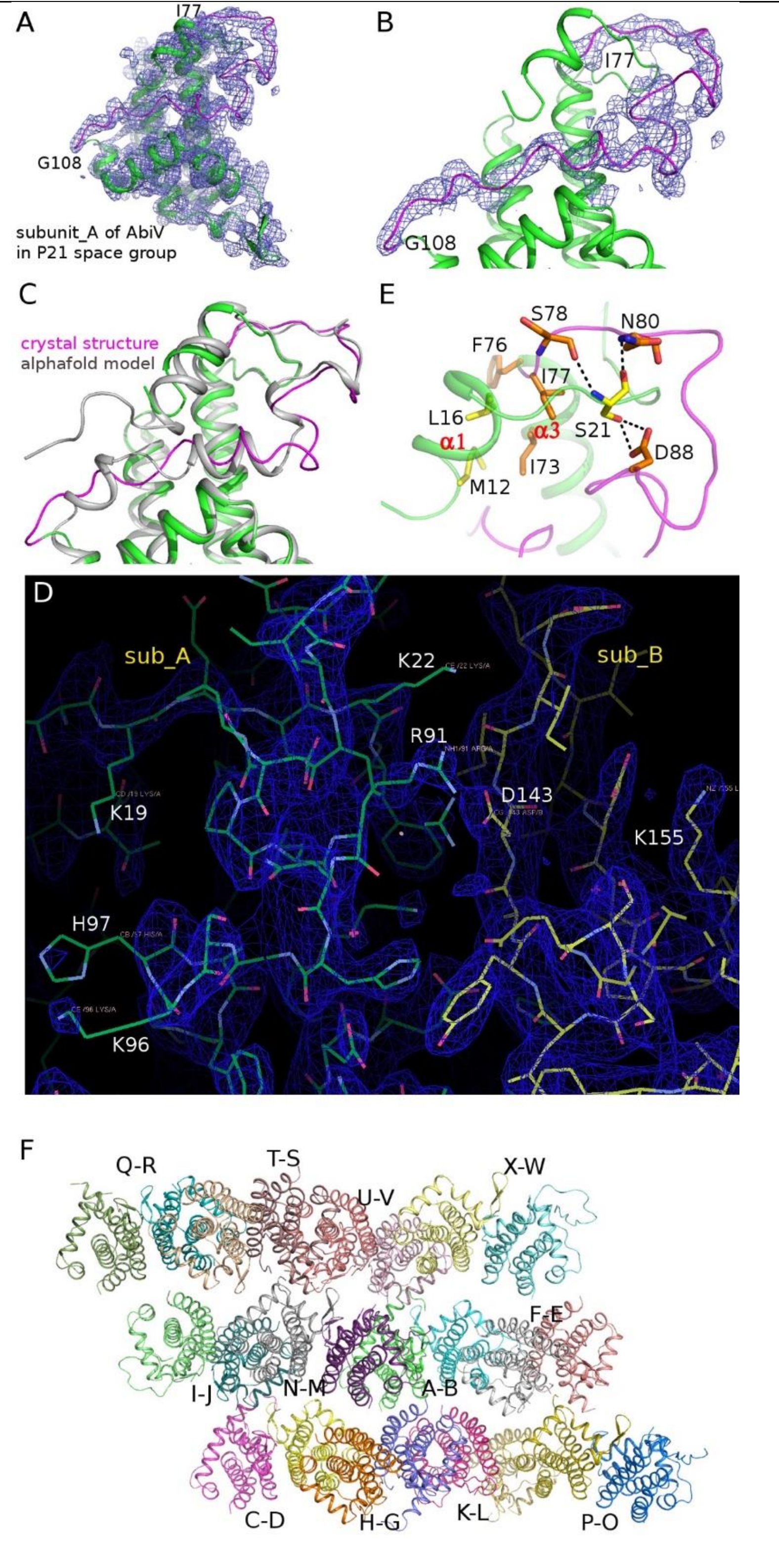


**Supplementary Figure 2.** Crystal structures of AbiV. Panel A) The overall electron density map (2Fo-Fc, contoured at 1σ) for the subunit A of AbiV in the monoclinic (P21) crystal form is shown in light blue. Cartoon representation of AbiV is shown in green with its flexible region (I77-G108) colored in magenta. Panel B) Close-up view of panel A showing the electron density map for the region I77-G108. Panel C) Superposition of alphafold model (gray) of AbiV onto its crystal structure (green and magenta) indicating the significant deviation in the conformation of the flexible region (magenta). Panel D) The relative positioning of R91 and H97 in the $RX_{4-6}H$ motif in the subunit A of AbiV as shown from a Coot screenshot. The electron density map (2Fo-Fc, blue) is contoured at 1σ. The clearly defined side chain of R91 forms H-bonds with D143 of subunit B while the electron density for the H97 side chain is very weak. Notably, the side chains of many lysine residues (e.g., K19, K22, K96) on the surface are not well defined although this crystal was obtained after the surface lysine methylation of AbiV. Panel E) Representative interactions between the N-terminal α1-helix and the α3-helix as well as the following loop α3/α4, exemplified by the van der Waals contacts between M12 and L16 in α1 and I73, F76, and I77 in α3 as well as by the H-bonds formed between S21 in α1 and S78, N80, and D88 in the loop α3/α4. Panel F) Cartoon representations of 24 AbiV_Δ23 subunits (subA to subX) forming 12 dimers in the asymmetric unit of the AbiV_Δ23 crystal.

**Supplementary Table 1**: Bacterial stains, phages, plasmids, and oligonucleotides.

| Materials | Relevant characteristics | Source |
|---|---|---|
| **Bacteria** | | |
| *E. coli* | *E. coli* M15; $Amp^R$; $Cam^R$ | 1 |
| *L. cremoris* $AbiV^S$ | *L. cremoris* MG1363; $Cam^R$ | This study |
| *L. cremoris* $AbiV^R$ | *L. cremoris* MG1363 transformed with pNZ123, carrying AbiV system; $Cam^R$ | This study |
| **Phages** | | |
| p2 | *Skunavirus*, infecting *L. cremoris* MG1363 | FHRCBV |
| c2 | *Ceduovirus*, infecting *L. cremoris* MG1363 | FHRCBV |
| **Plasmids** | | |
| pNZ123 | High copy number, $Cam^R$, 2.5kb | 2 |
| FL_*abiV*::pNZ123 | pNZ123 carrying full-length AbiV system | This study |
| *abiV1*::pNZ123 | pNZ123 carrying *abiV1* | This study |
| Y8X::pNZ123 | pNZ123 carrying full-length AbiV with a stop codon substitution at Y8 in *abiV2* | This study |
| AbiV::pQE70 | pQE70 carrying protein AbiV | 1 |
| AbiV_Δ14::pQE70 | pQE70 carrying truncated AbiV without 14 aa at N-terminus | This study |
| AbiV_Δ23::pQE70 | pQE70 carrying truncated AbiV without 23 aa at N-terminus | This study |
| AbiV_R91A::pQE70 | pQE70 carrying AbiV with R91A single substitution | This study |
| AbiV_R91A/H97A::pQE70 | pQE70 carrying AbiV with R91A and H97A dual substitutions | This study |
| **Oligonucleotides** | **Sequence 5' to 3'** | |
| FL_*abiV*::pNZ123_Fw | ATTACAGCTCCAGATCCAGTACTGAATTCTAAAAAGAGAGTGGGTGTATC | This study |
| FL_*abiV*::pNZ123_Rv | GAAAATATGCACTCGAGAAGCTTGAGCTCTGAATAGTGTTGATGAATTTTATC | This study |
| *abiV1*::pNZ123_Fw | ATTACAGCTCCAGATCCAGTACTGAATTCTACGGAGAGTTTTATGTTTGATAAAG | This study |
| *abiV1*::pNZ123_Rv | GAAAATATGCACTCGAGAAGCTTGAGCTCTCAGAATATTCAGCAAAATCAAACTC | This study |
| Y8X::pNZ123_Fw | GACAAAACTTAGTCTAAAATTGATAC | This study |
| Y8X::pNZ123_Rv | TATTTCATCCAAAAAGTCTTTGTTTTC | This study |
| wR19X::pNZ123_Fw | CAATACTAAAGTTTGACAAACTGAAG | This study |
| R19X::pNZ123_Rv | ACAGTATCAATTTTAGAGTAAGTTTTG | This study |
| AbiV_Δ14::pQE70_Fw | CTTAATACCAAAGAAAGTAAGTTTTC | This study |
| AbiV_Δ14::pQE70_Rv | CATGCTTAATTTCTCCTCTT | This study |
| AbiV_Δ23::pQE70_Fw | CTAAAGTCAACTGATGATCT | This study |

| | | |
|---|---|---|
| AbiV_Δ23::pQE70_Rv | CATGCTTAATTTCTCCTCTT | This study |
| 17-mer short RNA | AAUUGAUACUGUCAAUA | This study |
| 18-mer short RNA | AAGUAGCUGCAACUACUA | This study |
| Transcript of *abiV2* | AUAAAGAAAACAAAGACUUUUUGGAUGAAAUAGACAAAACUUACUCUAAAAUUGAUACUGUC<br>AAUACUAAAGUUAGACAAACUGAAGUAGCUGCAACUACUAAUCAACUUGCGCUAACUAAAGC<br>AAAUGUACAAAUUCAUACCCUUUUAGUAAUUGCUAGUAAUUAUUAUCAAUCAGUAUGGAUCC<br>AGAUUAAAGAAUGAACGGAGAGUUUUAUGUUUGAUAAAGACAACUAUGCAUUAGGAAAAAUG<br>AAGAAUACCCUUAAUACCAAAGAAAGUAAGUUUUCUCUAAAGUCAACU | This study |
| **Deposited data** | | |
| Structure of AbiV | 9BJ5, 9BJ6 | This study |
| Structure of AbiV_Δ23 | 9BJ7 | This study |

AbiV$^S$: AbiV phage sensitive phenotype; AbiV$^R$: AbiV phage resistance phenotype; Amp$^R$: ampicillin resistance (100 µg ml$^{-1}$); Cam$^R$: chloramphenicol resistance (5 µg ml$^{-1}$); FHRCBV, Félix d'Hérelle Reference Center for Bacterial Viruses.

**Supplementary Table 2**: Probable transcription start sites and terminators on FL_abiV::pNZ123

| Positions of Putative TSS | Coverage | Remarks |
| --- | --- | --- |
| 469 | 1212936 | + sense |
| 710 | 9862 | - sense |
| 1029 | 1070 | - sense |
| 1231 | 246 | + sense |
| 2256 | 128 | - sense |
| 2431 | 91 | - sense |
| **2558** | **289** | **+ sense** |
| **2617** | **629** | **+ sense** |
| **3222** | **813** | **+ sense** |
| **3286** | **671** | **+ sense** |
| 3521 | 973 | - sense |
| **Positions of Putative TTS** | **Coverage** | **Remarks** |
| 661 | 119025 | + sense |
| **2254** | **83747** | **+ sense** |
| 2577 | 704 | - sense, high background noise |
| **2853** | **414** | **+ sense, high background noise** |
| **3553** | **2239** | **+ sense** |
| 3747 | 766 | + sense |
